# Location of maximum bending stress in tapered hollow plants: A case study on Moso bamboo (*Phyllostachys edulis*)

**DOI:** 10.64898/2026.11.16.767008

**Authors:** Carol Lee Chalermsin, Satoru Tsugawa, Motohiro Sato

## Abstract

Many slender plants, such as trees and bamboo, have naturally tapered, axially graded forms to withstand wind loads; however, while trees feature solid cross-sections, bamboo culms are hollow. Although tapered hollow stems improve mechanical efficiency, the location at which the maximum bending stress develops remains unclear. As failure in such structures is driven by the bending-induced stress state, identifying this location is essential for understanding their structural capacities. This study aimed to (1) identify the location of the maximum bending stress of a tapered hollow beam made from an axially functionally graded material under various load profiles, (2) verify the consistency between linear and large deflection analyses, (3) investigate the effect of taper ratio on this location, and (4) apply the analysis to Moso bamboo (*Phyllostachys edulis*) as a case study. Governing nonlinear equations were derived for uniformly distributed load, inverse triangular load, triangular load, and point load; the equations were solved using a large deflection analysis. A linear analysis was also conducted and compared with the nonlinear results. The results showed that the linear analysis is sufficient for predicting the location of the maximum stress under distributed loading; however, under concentrated loading, it is accurate only at low loads. A more tapered geometry shifted the maximum stress toward the tip. In Moso bamboo, the maximum stress is located in the upper region of the culms, providing insights into the natural structural design of tapered hollow plants and a foundation for the design of bio-inspired tapered hollow components.

**Key Message:** The location of the maximum bending stress depends on load type and taper ratio; below ratios of 1/3−2/3, it shifts upward, reaching 83−96% of culm height in Moso bamboo.

## 1. Introduction

Many slender plants are tapered, narrowing from the base to the tip (Inoue et al. 2021; Kanahama and Sato 2023); however, they differ in cross-section. Trees are typically solid, whereas bamboo is hollow (Kanahama and Sato 2023). The hollow, slender form of bamboo is particularly intriguing because it is counterintuitive. On the surface, such a thin-walled tubular structure appears fragile, seems incapable of surviving harsh environments, and is prone to cross-sectional flattening (ovalization) under bending. Nevertheless, bamboo can withstand severe and repeated wind loads throughout its lifetime, suggesting that its apparent fragility conceals an efficient underlying mechanical design. The natural structure of bamboo is tapered and hollow with nodes and internodes. The vascular bundles in the structure grow radially from the inner wall to the outer wall, and their average volume fraction increases with height (Amada et al. 1997). These features make bamboo an axially and radially functionally graded material (FGM). Bamboo is a fascinating plant, and its nodes and vascular bundles are strategically arranged to strengthen the culm, minimize ovalization, and enhance resistance to bending (Chalermsin et al. 2025; Sato et al. 2017; Shima et al. 2016).

Previous studies have analyzed the mechanical behavior of tapered plant structures under various loading conditions. Horibe and Mori (2018) derived a bending stress equation for a solid, tapered, axially functionally graded beam; Nishiyama and Sato (2022) considered a tapered hollow beam under a concentrated tip load to obtain the deflection and bending stress through a large deflection analysis. A concentrated load approximates a sheltered, forest-grown plant shielded by its neighbors. However, for an unsheltered or open-grown plant, wind load is better represented as distributed along the length. Chalermsin and Sato (2025) addressed this loading scenario by analyzing a solid and hollow tapered axially functionally graded beam with a uniformly distributed load (UDL), obtaining deflection curves and normalizing the bending stress (from a linear analysis). However, real environmental loading varies with height and is not uniform. Therefore, a UDL alone cannot capture the range of load profiles that a culm or trunk may experience.

Although Horibe and Mori (2018) and Chalermsin and Sato (2025) identified the threshold taper ratios at which the location of maximum stress shifts under a point load and UDL, respectively, it remains unclear how hollow cross-section and load magnitude affect the location of maximum stress in a tapered hollow axially functionally graded beam under various load profiles. In addition, the results from a linear analysis have not been directly compared with those from a large deflection analysis, making it unclear how geometric nonlinearity influences the location of the maximum stress. Identifying the location of maximum bending stress for different structural designs and loading conditions is essential for predicting where failure is most likely to occur, which was the motivation behind our work.

The objectives of this study were: (1) to identify the location of the maximum bending stress of a tapered hollow beam made from an axially FGM under various load profiles, (2) to verify the consistency between linear and large deflection analyses, (3) to investigate the effect of taper ratio on the location of maximum stress under different loading conditions, and (4) to apply the analysis to Moso bamboo (*Phyllostachys edulis*) as a case study of a naturally occurring tapered hollow structure. Four load profiles were considered: UDL, inverse triangular load, triangular load, and point load. Although real wind loading does not strictly follow these idealized profiles, the analysis clarifies the influence of load distribution on the location of the maximum bending stress.

To investigate this relationship, we modeled trees or bamboo as beams and analyzed the large deflection of solid and hollow tapered axially functionally graded beams under various load types. The governing equations for the beam were derived using the Euler–Bernoulli principle. To consider the axially functionally graded nature of bamboo, the Young’s modulus is assumed to vary linearly and is calculated as the Young’s modulus ratio. Governing equations for beams under different loads were derived. The governing equation for a beam subjected to UDL was derived using the method proposed by Chalermsin and Sato (2025). After obtaining the governing equations, we derived a dimensionless bending stress equation. The fourth-order Runge–Kutta method was used to solve the governing equations. A linear analysis was also conducted to derive the stress ratio equation and determine the location of maximum stress. The results were compared with those obtained under point loading.

## 2. Methodology

### 2.1 Models and Methods

This study considered solid and hollow tapered beams under various loading conditions and compared them with a point-load case. **Figure 1** shows a diagram of a bending tree and bamboo modeled as a solid tapered beam under UDL *q*, inverse triangular load *N*, triangular load *T*, and point load *P*; the Young’s modulus, diameter, and second moment of area at the base are denoted by *E*_0_, *D*_0_, and *I*_0_, and those at the tips are denoted by *E*_1_, *D*_1_, and *I*_1_, respectively. *L, D*(*s*), and *s* denote the total length, linearly varying diameter, and distance from the base to the point along the length, respectively. **Figure 1(a)** shows an unsheltered or open-grown bending tree modeled as a solid tapered beam subjected to a distributed load; other than uprooting, the other common failure mode of bamboo is trunk breakage (**Fig. 1(b)**); **Figures 1(c), (d)**, and **(e)** correspond to realistic wind distributions. As for sheltered or forest-grown plants, where surrounding plants provide shelter, such as bamboo, as shown in **Fig. 1(e)**, the dominant failure mode of hollow trunks is longitudinal splitting (Huang et al. 2017); solid tapered beam under a point load (**Fig. 1(f)**) is modeled. Under bending, stress develops in the culm, and the location of the maximum stress is prone to failure. To investigate this, the conditions presented in **Figs. 1(c), (d), (e)** and **(h)** were modeled.

**Figure 1.**
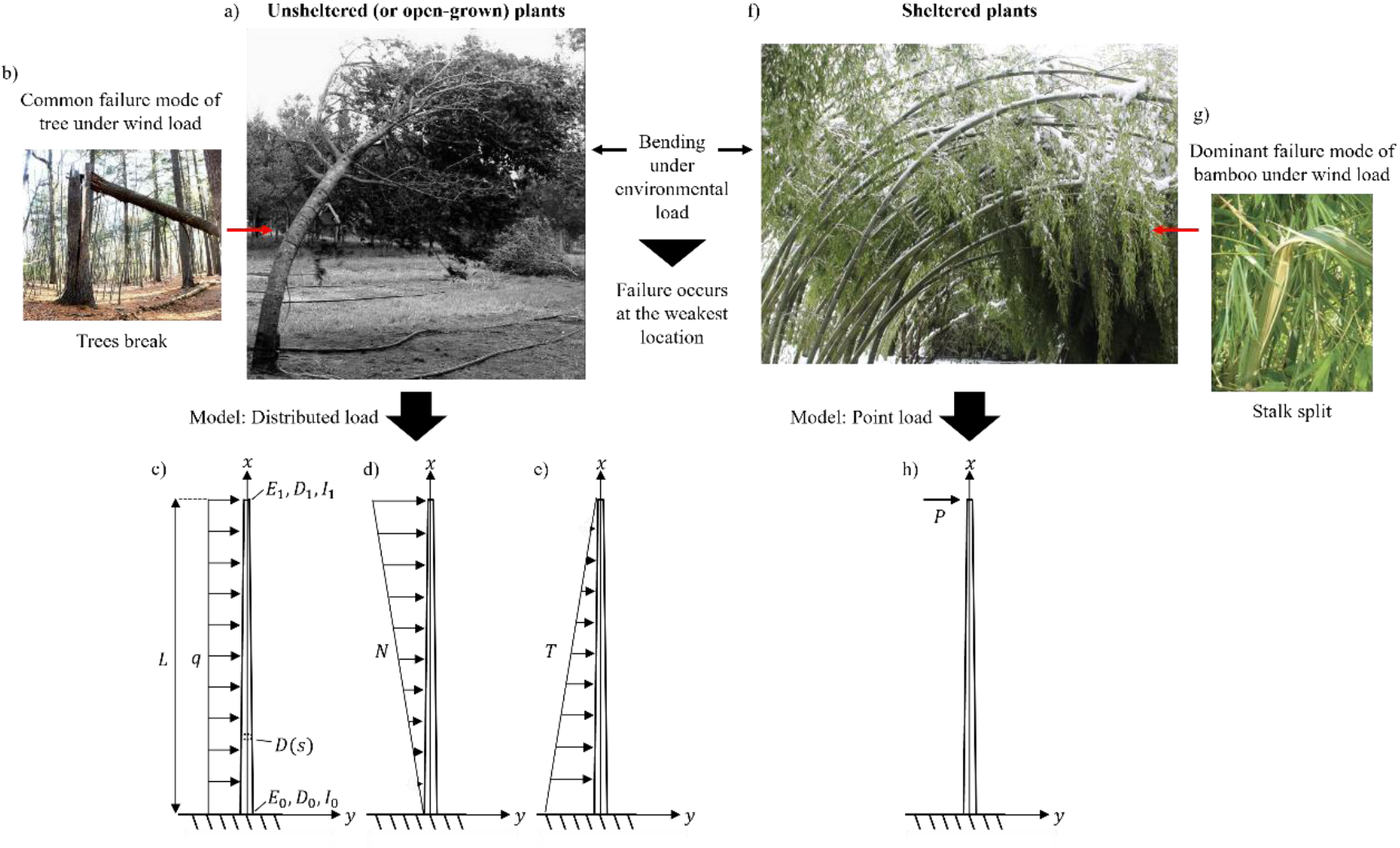
Solid tapered beam subjected to **(a)** Bending tree [reproduced with permission from Gilman et al. (2008), © International Society of Arboriculture], **(b)** Conventional bending failure of trees under wind load [Photograph courtesy of A. Ponomarenko, reproduced with permission (Randall 2016)], **(c)** Uniformly distributed load *q*, **(d)** Inverse triangular load *N*, **(e)** Triangular load *T*, **(f)** Bending bamboo [Photograph © Hans Friederich, reproduced with permission (Friederich 2021)], **(g)** Longitudinal splitting is the dominant failure mode of bamboo under wind load [reproduced from Huang et al. (2017) under CC BY 4.0], **(h)** Point load *P*.

In this study, the diameter was assumed to vary linearly with *D*(*s*) (Amada et al. 1997). As the Young’s modulus of bamboo increases with height (Amada et al. 1996; Horibe and Mori 2018; Kamruzzaman et al. 2008), this study assumed a linearly varying Young’s modulus *E* and moment of inertia *I*. The flexural rigidity can be calculated using the following equation:

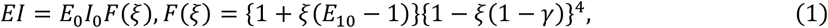

where *E*_10_ = *E*_1_/*E*_0_ is the Young’s modulus ratio.

Assuming that the volume of the solid untapered rod 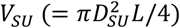 and tapered rod 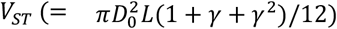 are equal (*V*_*ST*_ = *V*_*SU*_), where *D*_*SU*_ is the diameter of the solid untapered beam, the following relationships can be obtained:

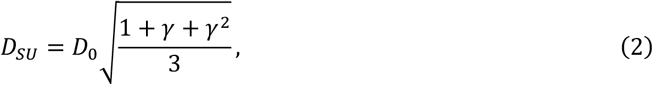

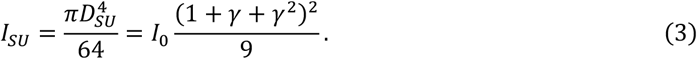

#### 2.1.1 Case of uniformly distributed load (UDL)

Several experimental studies have used the Euler–Bernoulli beam theory to analyze the bending of bamboo culms (Chalermsin and Sato 2025; Horibe and Mori 2018; Mouka et al. 2022; Nishiyama and Sato 2022). Therefore, this theory was used in this study.

The governing equation for solid tapered beams under UDL was derived using a similar procedure as employed by Chalermsin and Sato (2025); however, the dimensionless load *α*_*q*_ was used instead, expressed as *α*_*q*_ = *qL*^3^/*E*_*SU*_ *I*_*SU*_, where *E*_*SU*_ is the Young’s modulus, and *I*_*SU*_ is the second moment of area of a solid untapered beam.

The governing equation for solid tapered beams under UDL can be described as follows:

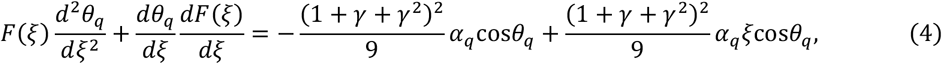

where *ξ* = *s*/*L, γ* = *D*_1_/*D*_0_ is the taper ratio, and *θ*_*q*_ is the slope. *F*(*ξ*) follows Eq. (1). By differentiating

*F*(*ξ*) with respect to *ξ*, we can obtain the following equation:

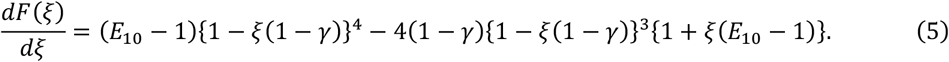

Using the method employed by Chalermsin and Sato (2025), the area of the hollow section *A*_*HT*_(*s*) and volume of the hollow tapered 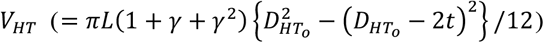 calculated. To derive the governing equation for the hollow tapered beam under the UDL, the volume of the solid tapered beam was set to be equal to that of the hollow tapered beam 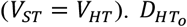 can be expressed using *D*_*o*_ as shown below (Chalermsin and Sato 2025):

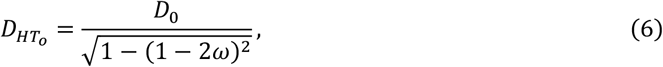

where 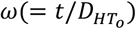 is the thickness divided by the outer diameter ratio (0 < *ω* ≤ 0.5).

By using the Euler–Bernoulli and dimensionless load, *α*_*q*_ = *qL*^3^/*E*_*SU*_ *I*_*SU*_ e an xpress the governing equation for hollow tapered beams under UDL as follows:

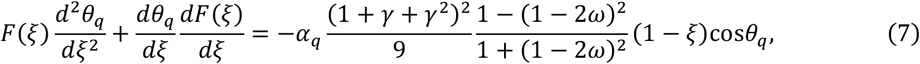

where *F*(*ξ*) and *dF*(*ξ*)/*dξ* follow Eqs. (1) and (5).

#### 2.1.2 Case of inverse triangular Load *N*

**Figure 2** shows a diagram of a solid tapered beam subjected to an inverse triangular load with the fixed end as the origin.

**Figure 2.**
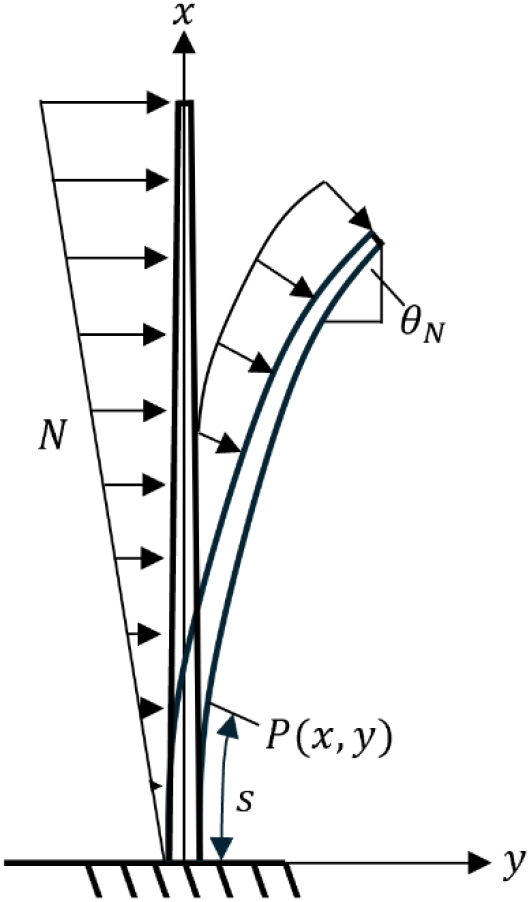
Solid tapered beam under inverse triangular load *N*; *θ*_*N*_ is the slope.

The Euler–Bernoulli equation for a beam under an inverse triangular load is as follows:

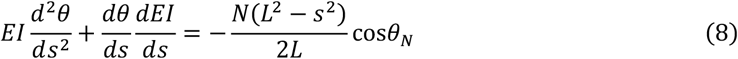

where *EI* follows Eq. (1), and *N* denotes an inverse triangular load. By substituting the dimensionless load *α*_*N*_ = *NL*^3^/*E*_*SU*_ *I*_*SU*_ in Eq. (8) and simplifying it, we can express the governing equation for a solid tapered beam under *N* as follows:

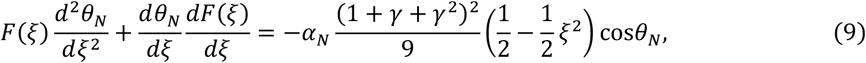

where *F*(*ξ*) and *dF*(*ξ*)/*dξ* follow Eqs. (1) and (5), respectively.

Using a procedure similar to that described in Section 2.1.1, but with *N*, the governing equation for the hollow tapered beam under *N* can be expressed as follows:

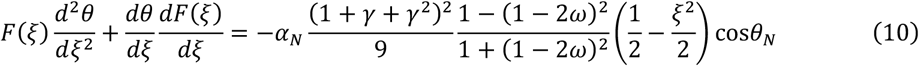

#### 2.1.3 Case of triangular load *T*

Using a similar procedure, we can express the governing equation for a solid tapered beam under triangular load *T* as follows:

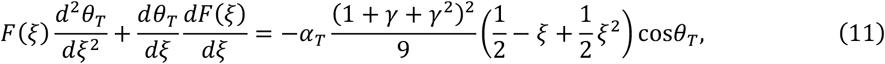

where *F*(*ξ*) and *dF*(*ξ*)/*dξ* follow Eqs. (1) and (5), respectively.

The corresponding governing equation for hollow tapered beams under *T* is:

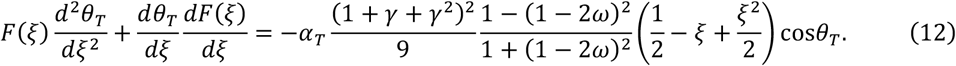

### 2.2 Derivation of dimensionless bending stress equation

After deriving the governing equations for the beam under the three distributed loads, the next task was to derive an equation for the dimensionless bending stress. We refer to the bending stress equation, *σ* = *Mr*/*I*, and the moment–curvature relationship, *M* = *EIdθ*/*ds*, where *M* is the bending moment, *r* is the radius, *E* is the Young’s modulus, *I* represents the second moment of area, *θ* is the slope, and *s* is the position along the deflected beam. Nishiyama and Sato (2022) derived a dimensionless bending stress equation for a solid tapered beam; however, this study extended it to hollow tapered axially functionally graded beams as follows:

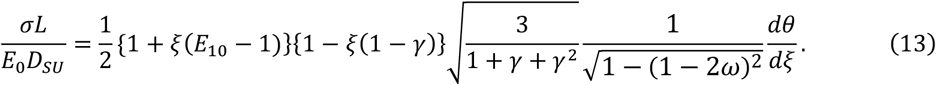

### 2.3 Linear analysis: Normalized bending stress

Using a linear analysis, we derived the normalized bending stress equation for a solid tapered beam under distributed loads, as well as the location of the maximum stress.

#### 2.3.1 Case of UDL

Let *σ*_*q*_(*x*) be the bending stress of a solid tapered beam, i.e., *σ*_*q*_(*x*) = *M*(*x*)*r*(*x*)/*I*(*x*) here *M*(*x*), *r*(*x*), and *I*(*x*) represent the bending moment, radius of the uniformly tapered beam, and second moment of area, respectively. *σ*_*q*0_ = 16*qL*^2^/*πD*^3^ represents the outer stress at the fixed end, and *D* represents the diameter of the straight homogeneous beam. For the UDL, the normalized bending stress can be calculated as follows (Chalermsin and Sato 2025):

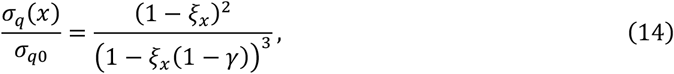

where *ξ*_*x*_ = *x*/*L* By differentiating *σ*_*q*_(*x*)/*σ*_*q*0_ with respect to *ξ*_*x*_ we can obtain the location of the maximum stress *ξ*_*qmax*_ using Eq. (15). At *ξ*_*qmax*_, the maximum stress (*σ*_*q*_(*x*)/*σ*_*q*0_)_*max*_ can be calculated using Eq. (16).

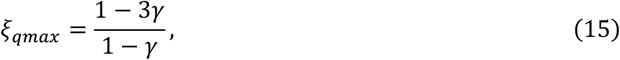

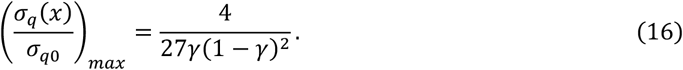

#### 2.3.2 Case of inverse triangular load *N*

By using a similar procedure as that employed for the UDL case, we can calculate the bending stress of a solid tapered beam under inverse triangular load *N* using *σ*_*N*_(*x*) = *M*(*x*)*r*(*x*)/*I*(*x*). The normalized bending stress equation for the beam under *N* is:

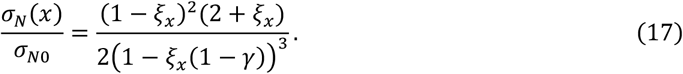

By differentiating Eq. (17) with respect to *ξ*_*x*_, we can obtain *ξ*_*Nmax*_, and at *ξ*_*Nmax*_, the maximum stress ratio (*σ*_*N*_(*x*)/*σ*_*N*0_)_*Tmax*_ can be calculated using the expression:

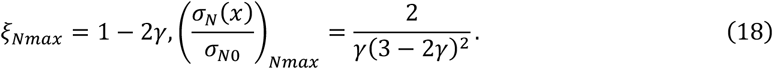

#### 2.3.3 Case of triangular load *T*

The normalized bending stress equation for beams under *T* is:

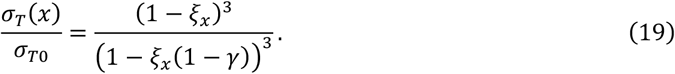

By differentiating Eq. (19) with respect to *ξ*_*x*_, we can obtain *ξ*_*Tmax*_, and at *ξ*_*Tmax*_, the maximum stress ratio (*σ*_*T*_(*x*)/*σ*_*T*0_)_*Tmax*_ can be calculated using the expression:

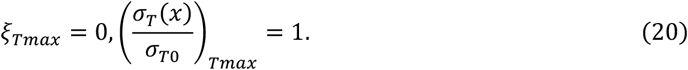

*ξ*_*Tmax*_ = 0 means that the location of maximum stress is always at the base, and (*σ*_*T*_(*x*)/*σ*_*T*0_)_*Tmax*_ = 1 indicates that the maximum normalized stress is 1.

To solve the governing equations for the beam under each loading condition, the Runge–Kutta and Regula–Falsi methods were used to obtain numerical solutions (Chalermsin and Sato 2025; Horibe and Mori 2018; Nishiyama and Sato 2022). The numerical program Mathematica software was used to solve these equations. The method and equations derived in this study were validated by calculating the deflection angles using the governing equation for the UDL (Chalermsin and Sato 2025). The method and equations were proven, as the calculated angles corresponded perfectly with the results reported by Xiao (2011) and Chalermsin and Sato (2025). Xiao (2011) investigated the large deflection of a beam subjected to UDL and obtained the deflection angles using the homotopy analysis method. Chalermsin and Sato (2025) analyzed the large deflection of hollow tapered axially functionally graded beams under a UDL.

## 3 Results and discussion

### 3.1 Systematic evaluation of normalized bending stress of beam subjected to different loading conditions

**Figures 3(a), (b), (c)**, and **(d)** show the results of the normalized bending stress of the beam under the UDL (Chalermsin and Sato 2025), inverse triangular load, triangular load, and point load (Horibe and Mori 2018), respectively, obtained using a linear analysis. For beams under the UDL, as shown in **Fig. 3(a)**, Chalermsin and Sato (2025) found that the location of maximum stress shifts away from the base when *γ* < 1/3 ≈ 0.33, and that this location and the corresponding maximum stress can be calculated using Eqs. (15) and (16), respectively. As shown in **Fig. 3(b)**, for the beam subjected to an inverse triangular load, the location of the maximum stress shifts away from the base when *γ* < 1/2. When the beam is subjected to a triangular load, as shown in **Fig. 3(c)**, the location of the maximum bending stress is at the base regardless of *γ*, which follows Eq. (20). **Fig. 3(d)** shows the normalized bending stress of the beam under a point load (Horibe and Mori 2018). According to Horibe and Mori (2018), the location of the maximum stress shifts away from the base when *γ* < 2/3 ≈ 0.67. In addition to the observation made by Chalermsin and Sato (2025) who concluded that the type of load affects *γ*, which causes the location of the maximum stress to shift, this result shows that for beams under inverse triangular load, the threshold of *γ* is different from that observed under the UDL and point load. As for the triangular load, the location of maximum stress is at the base regardless of *γ*.

**Figure 3.**
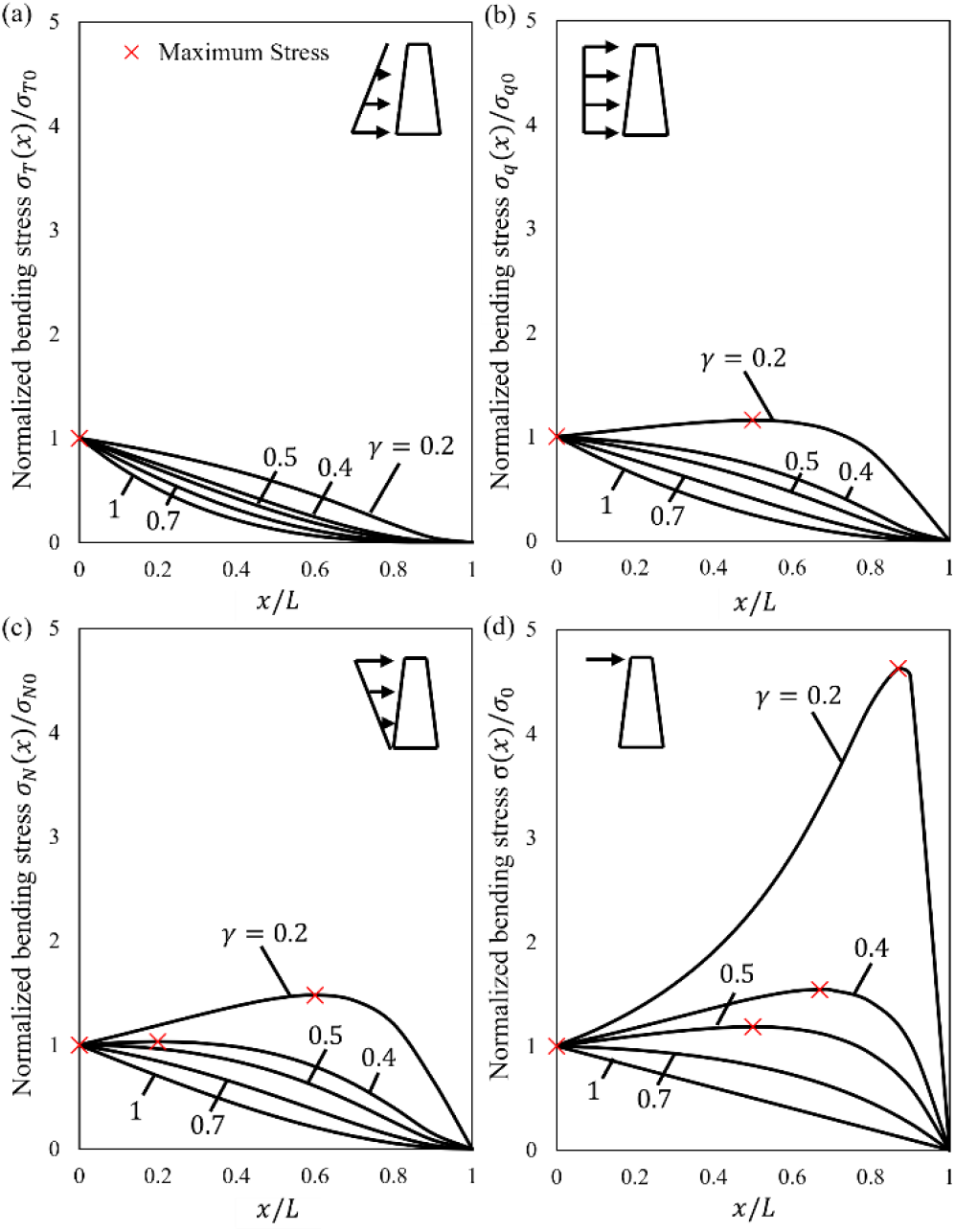
Normalized bending stress of beam under linear analysis: **(a)** Uniformly distributed load (UDL) (Chalermsin and Sato 2025), **(b)** Inverse triangular load, **(c)** Triangular load, and **(d)** Point load (Horibe and Mori 2018) with varying taper ratio *γ*.

These results and the equations derived are beneficial for structural mechanics and structural design, as they emphasize the importance of considering load types and *γ*.

Fiber bundles of bamboo that grow radially from the inner to the outer surface of the cross section (Javadian et al. 2019; Zhang et al. 2024) do not affect the location of maximum stress; however, such a graded structure decreases the stress distribution along the height (Sayyad et al. 2024). Sayyad et al. (2024) compared uniform and graded arrangements of fiber bundles in a cross-section and tapered beam form. The axial stress in the beam was investigated under a concentrated load applied to the free end. The locations of maximum stress for both the homogeneous and graded cases were the same, and the stress distribution along the height exhibited a similar trend; however, the stress level of the graded case was lower by approximately 75%.

### 3.2 Location of maximum stress under different loading conditions

**Figure 4** shows the location of maximum stress with varying *γ* under inverse triangular load, point load, and UDL. Under the triangular load, the maximum always occurred at the base; therefore, it is not plotted. Two morphological datasets were superimposed: 16 six-year-old Moso bamboo populations from China (Zhang et al. 2024) and a two-year-old culm from Japan (Amada et al. 1996, 1997). For the Chinese data, diameters were measured only below the first branch, and the taper ratio was evaluated as *γ*_*FB*_ = *D*_*FB*_ /*D*_0_ here *D*_0_ and *D*_*FB*_ are the maximum and minimum internode diameters reported for each population, giving *γ*_*FB*_ = 0.36 − 0.53. Since these extremes are taken from different culms within a population, the ratio underestimates the taper ratio of any individual culm and should be considered a lower bound; the actual *γ*_*FB*_ lies further to the right in **Fig. 4**.

Consequently, fewer populations intersected the load curves than those shown, and when no intersection occurred, the maximum stress was located at the base. The Japanese data (*γ* = 0.08) intersected all three curves close to the tip and covered the entire culm, making them the more representative case for the present analysis. For the branch-free segment described by *γ*_*FB*_, the point load is the most appropriate profile, since wind load acting on the crown is transmitted to this segment essentially as a concentrated force at the first branch.

**Figure 4.**
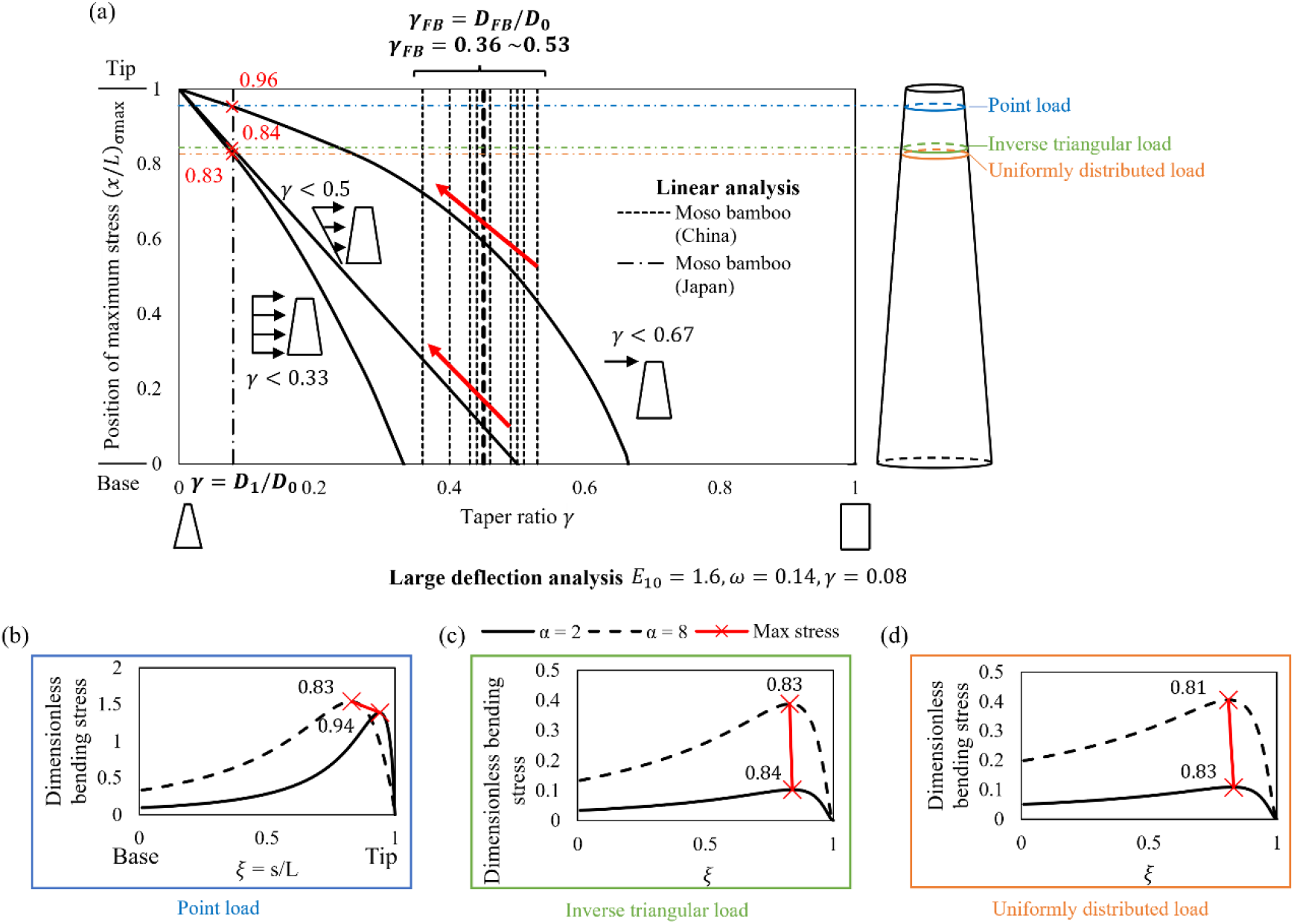
**(a)** Location of the maximum stress (*x*/*L*)_*σmax*_ against the taper ratio of the culm from the base up to the first branch *γ*_*FB*_ of a tapered beam under point, inverse triangular, and uniformly distributed loads (linear analysis) compared with morphological Moso bamboo data from China (Zhang et al. 2024) and taper ratio *γ* from Japan (Amada et al. 1997). *γ*_*FB*_ = *D*_*FB*_ /*D*_0_, where *D*_*FB*_ is the diameter of the internode under first branch and *D*_0_ is the diameter at the base. (b-d) Dimensionless bending stress from latge deflection analysis based on the morphological data from Amada et al. (1997) (*E*_10_ = 1.6, *ω* = 0.14, *γ* = 0.08) under **(b)** point load, **(c)** inverse triangular load, and **(d)** uniformly distributed load, for *α* = 2 (solid lines) and *α* = 8 (dashed lines); × marks the maximum stress

Since *ω* and *E*_10_ of Moso bamboo vary with height, representative values of *γ* = 0.08, *ω* = 0.14 (average), and *E*_10_ = 1.6 were adopted from Amada et al. (1996, 1997). The insets in **Fig. 4** show the corresponding large deflection stress distributions under point load, inverse triangular load, and UDL for *α* = 2 and 8. Under point load, the location of maximum stress *ξ*_*σmax*_ = 0.94 at *α* = 2, which is in close agreement with the linear analysis ((*x*/*L*)_*σmax*_ ≈ 0.96), and it shifts toward the base to *ξ*_*σmax*_ ≈ 0.83 at *α* = 8. indicating peak migration under increasing load. Under distributed loads, the location of the maximum stress is nearly independent of the load magnitude: *ξ*_*σmax*_ = 0.84 and 0.83 under inverse triangular load, and 0.83 and 0.81 under UDL, for *α* = 2 and 8, respectively. Both these values are in agreement with the linear predictions (0.84 and 0.83). The linear and large deflection analyses are therefore consistent at low loads and deviate only under point loads with increasing *α*.

To isolate the effect of axial grading, the analysis was repeated for a homogeneous beam (*E*_10_ = 1) with the same *γ* and *ω* Under the UDL, both the location and the magnitude of the maximum stress remained essentially unchanged. Under inverse triangular loading, the results were also nearly identical, apart from the slight shift in *ξ*_*σmax*_ from 0.83 (*E*_10_ = 1.6) to 0.82 (*E*_10_ = 1) at *α* = 8 By contrast, under a point load, grading had a pronounced effect. The peak of the homogeneous beam lay closer to the base (*ξ*_*σmax*_ = 0.92 and 0.77 at *α* = 2 and 8, versus 0.94 and 0.83 for *E*_10_ = 1.6), and its maximum stress was approximately 27% lower (1.01 (*E*_10_ = 1) versus 1.39 (*E*_10_ = 1.6) at *α* = 2 1.13 (*E*_10_ = 1) versus 1.54 (*E*_10_ = 1.6) at *α* = 8).

### 3.3 Verification of linear and large deflection analysis and comparison with Moso bamboo morphological data

**Figure 4** shows that as *γ*_*FB*_ decreases (i.e., as the beam becomes more tapered), (*x*/*L*)_*σmax*_ shifts toward the tip. Moso bamboo with *γ*_*FB*_ = 0.53 intersects the point load curve at (*x*/*L*)_*σmax*_ ≈ 0.44, while *γ*_*FB*_ = 0.36 gives (*x*/*L*)_*σmax*_ ≈ 0.72, as indicated by the red arrows. Although *γ*_*FB*_ does not consider the entire culm, considering that all Moso bamboo species follow a linear trend, *γ*_*FB*_ is useful for understanding how tapering affects the shift in the stress, though the actual taper ratio *γ* of Chinese Moso bamboo would shift to the left, closer to the Japanese Moso bamboo.

The location of maximum stress differed substantially between the load types for the same taper ratio. Referring back to the bending stress equation *σ* = *My*/*I*, since *y* and *I* are the same regardless of the applied load, the stress distribution differs between load types only through the bending moment distribution *M*(*ξ*) Under point loading, *M*(*ξ*) decreases linearly from the base to the tip [*M*(*ξ*) = *P* · *L*(1 − *ξ*)]. Under inverse triangular loading, the moment distribution [*M*(*ξ*) = (*N*/*L*) · (*L*^3^/3 − *ξL*^3^/2 + *ξ*^3^*L*^3^/6)] decreases more rapidly, concentrating the moment closer to the base. Combined with the decrease in *I* toward the thinner tip, the slower reduction in the point load moment causes the maximum stress to occur in the upper region, whereas the more rapid moment decay of the inverse triangular loading places the maximum stress in the lower region (in comparison to the point load).

### 3.4 Case study on Moso bamboo: Location of maximum stress and safety factor under different loading conditions

**Tables 1(a)** and **(b)** summarize the location of the maximum stress (from both the linear and nonlinear analyses), corresponding maximum stress, location of the minimum safety factor *SF*_*min*_, and the safety factor for each load type under equal total forces of **(a)** 31.4 N and **(b)** 125.8 N. The safety factor *SF* is calculated by dividing the ultimate tensile strength with the maximum bending stress (*σ*_*ult*_/*σ*_*max*_) with *σ*_*max*_ obtained by converting the dimensionless bending stress to the physical bending stress. All the bamboo parameters were taken from Amada et al. (1996, 1997), who conducted experiments on two-year-old Moso bamboo (*E*_10_ = 1.6, *ω* = 0.14, *γ* = 0.08, *L* = 16 m, and *E*_0_ = 11 GPa) and reported a linearly increasing ultimate tensile strength from 140 MPa at the base to 235 MPa at the tip.

**Table 1.**
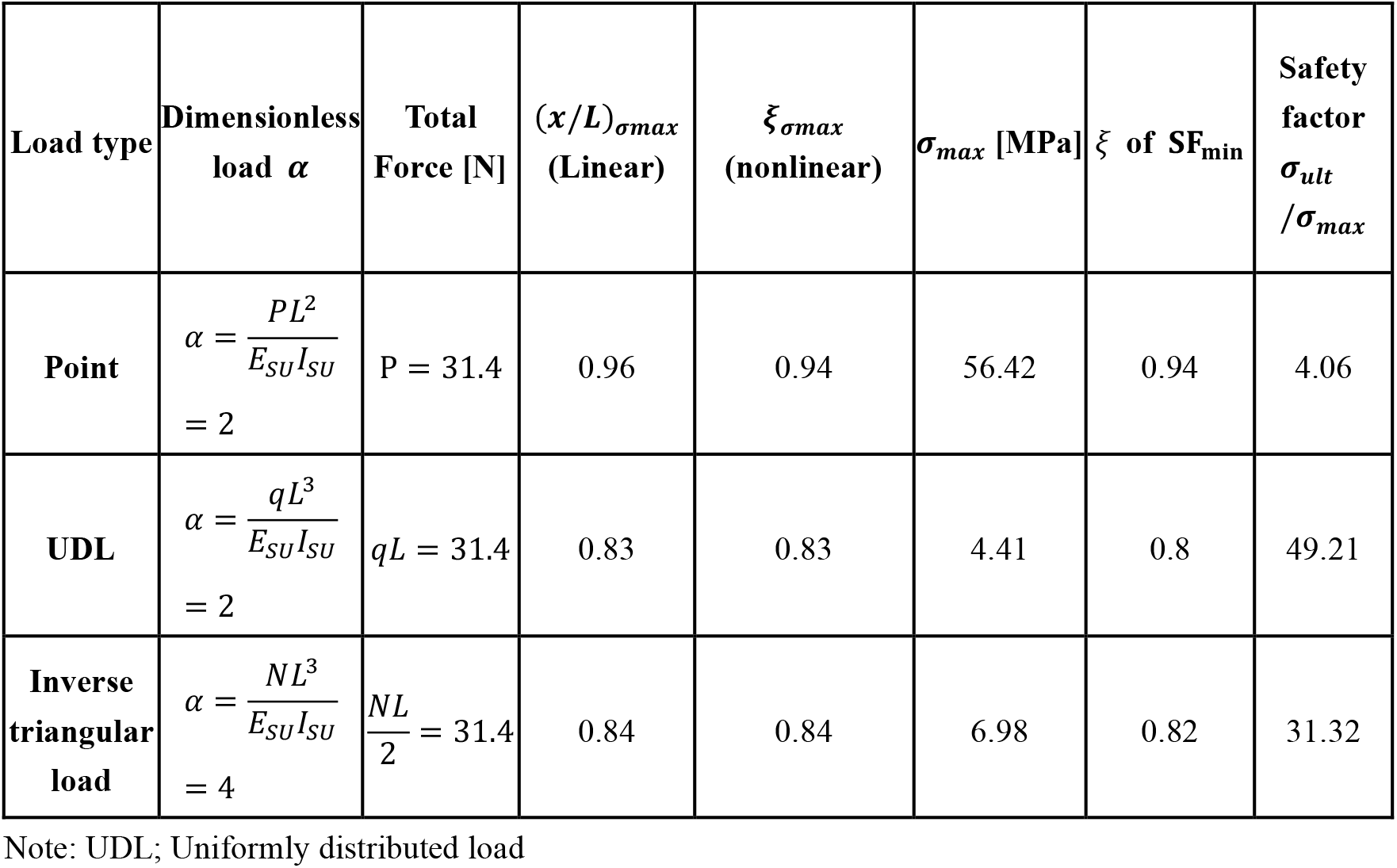
(a) Normalized location of maximum stress and safety factor of Moso bamboo grown in Japan (Amada et al. 1997): set total force = 31.4 [N].

| **Load type** | **Dimensionless load *α*** | **Total Force [N]** | **(*x*/*L*)<sub>*σmax*</sub> (Linear)** | ***ξ*<sub>*σmax*</sub> (nonlinear)** | ***σ*<sub>*max*</sub> [MPa]** | *ξ* **of SF<sub>min</sub>** | **Safety factor *σ*<sub>*ult*</sub> /*σ*<sub>*max*</sub>** |
|---|---|---|---|---|---|---|---|
| **Point** | $\alpha = \frac{PL^{2}}{E_{SU}I_{SU}} = 2$ | P = 31.4 | 0.96 | 0.94 | 56.42 | 0.94 | 4.06 |
| **UDL** | $\alpha = \frac{qL^{3}}{E_{SU}I_{SU}} = 2$ | *qL* = 31.4 | 0.83 | 0.83 | 4.41 | 0.8 | 49.21 |
| **Inverse triangular load** | $\alpha = \frac{NL^{3}}{E_{SU}I_{SU}} = 4$ | $\frac{NL}{2} = 31.4$ | 0.84 | 0.84 | 6.98 | 0.82 | 31.32 |

Note: UDL; Uniformly distributed load

**Table 1.**
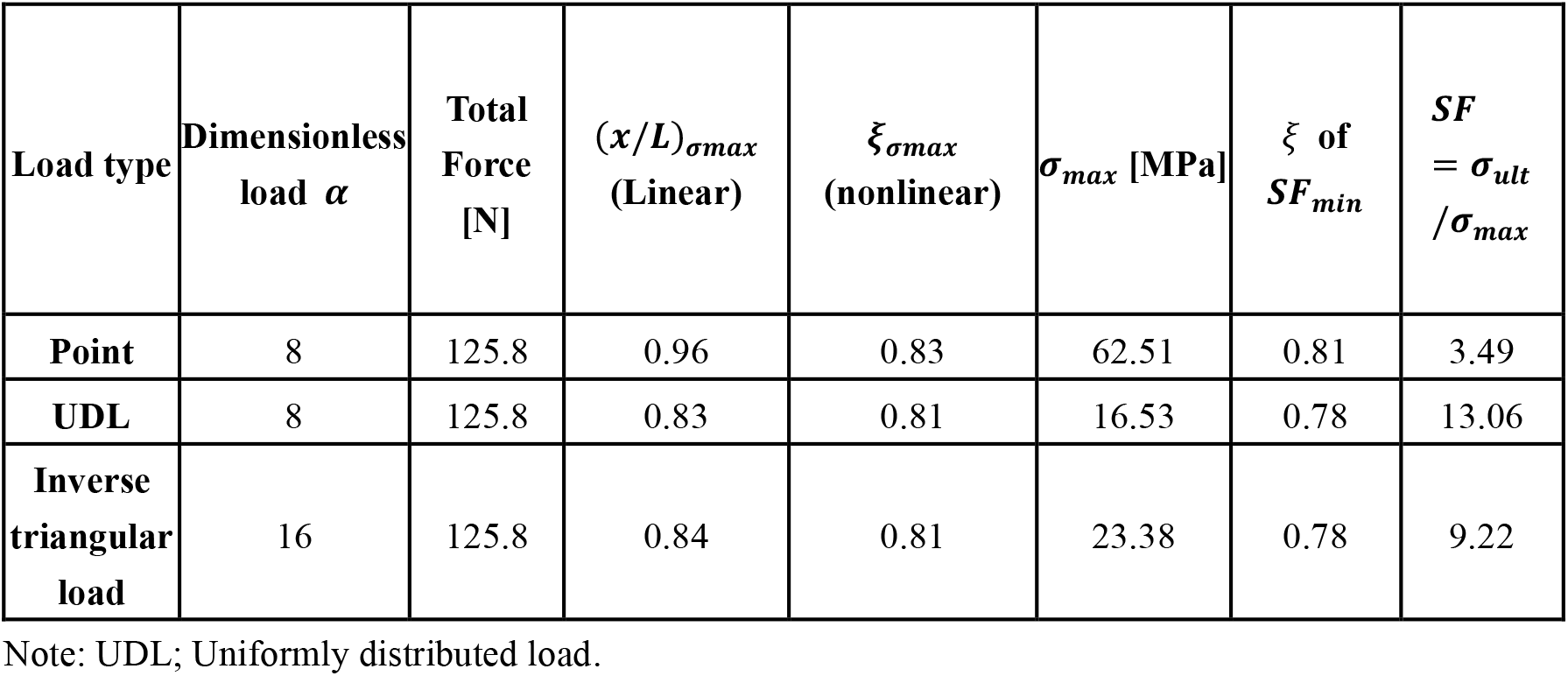
(b) Normalized location of the maximum stress and safety factor of Moso bamboo grown in Japan (Amada et al. 1996, 1997): set total force = 125.8 [N].

| **Load type** | **Dimensionless load** *α* | **Total Force [N]** | (*x*/*L*)<sub>*σmax*</sub> **(Linear)** | *ξ*<sub>*σmax*</sub> **(nonlinear)** | *σ*<sub>*max*</sub> **[MPa]** | *ξ* **of** *SF*<sub>*min*</sub> | *SF* = *σ*<sub>*ult*</sub> /*σ*<sub>*max*</sub> |
|---|---|---|---|---|---|---|---|
| **Point** | 8 | 125.8 | 0.96 | 0.83 | 62.51 | 0.81 | 3.49 |
| **UDL** | 8 | 125.8 | 0.83 | 0.81 | 16.53 | 0.78 | 13.06 |
| **Inverse triangular load** | 16 | 125.8 | 0.84 | 0.81 | 23.38 | 0.78 | 9.22 |

Note: UDL; Uniformly distributed load.

The location of maximum stress predicted by the linear and large deflection analyses were similar under low load (*α* = 2) but diverged with increasing load. At *α* = 8, they remained close for the UDL and inverse triangular load, but differed significantly for point loads, decreasing from 0.96 (linear) to 0.83 (large deflection). In all cases, the location of the maximum stress corresponded closely to the location of the minimum safety factor. These results confirm that the linear analysis is sufficient for determining the location of the maximum stress of a hollow, tapered, axially functionally graded beam under low loads and provides a quick estimation under high loads. Note that the linear (*x*/*L*)_*σmax*_ is identical in **Tables 1(a)** and **1(b)**, because it is independent of the load magnitude.

The location of the maximum stress (*ξ*_*σmax*_) of Moso bamboo lies away from the base and differs depending on the taper ratio and load type. From **Table 1**, for Japanese Moso bamboo (*γ* = 0.08) under point load, UDL, and inverse triangular load, *ξ*_*σmax*_ ranges from approximately *ξ* = 0.8 to 0.9 and corresponds closely with *ξ* of *SF*_*min*_ This shows that regardless of the load type, the location of the maximum stress of the Moso bamboo was close to the tip.

This study identified the location of the maximum stress and the influence of tapering and load types of a hollow tapered beam under various loading conditions, providing a foundation that could be extended into a design framework for engineering applications. Identification of the location of the maximum bending stress can enable a targeted reinforcement of tapered hollow and possible axial FGM structures, such as electric poles and composite tubes, in the construction and aerospace sectors.

This study has several limitations. The predicted locations of the maximum stress were theoretical, and experimental verification was required before application. This study used the ultimate tensile strength reported by Amada et al. (1996). However, longitudinal splitting is known to be the dominant failure mode in bamboo, meaning that actual failure may occur below the predicted axial tensile limit. Consequently, the safety factors reported in **Tables 1(a)** and **(b)** likely overestimated the true safety factor margin. This bio-inspired analysis simplifies real bamboo geometry; real culms have branches and nodes, which were neglected, and the hollowing ratio in practice decreases with height, contrary to the constant height assumed in this study.

## 4. Conclusions

This study analyzed the location of the maximum bending stress of a hollow, tapered, and axially functionally graded beam and examined how hollow and tapered geometries affect bending stress distribution under various loading conditions. Dimensionless and normalized bending stresses and nonlinear governing equations for solid and hollow tapered beams under various load types were derived and compared with practical data on Moso bamboo.

The four main objectives and corresponding results are as follows: (1) Various equations were derived to identify the location of maximum stress. For very tapered structures, such as the Japanese Moso bamboo, the location of the maximum stress lied consistently in the upper half of the culm for the load types considered. (2) The locations of the maximum stress obtained from the linear and large-deflection analyses were computed and compared. The linear analysis was found to be sufficient for locating the maximum stress under distributed loading, whereas under concentrated loading, it was accurate only at low loads. (3) The taper ratio caused the location of the maximum stress to shift along the height. The more tapered the beam, the more it shifted toward the tip. (4) By examining the Japanese Moso bamboo, we found that the location of maximum stress lies in the upper region of the culm.

These findings advance our understanding of natural structural design in plants, such as bamboo, and provide a foundation for the structural design of bio-inspired tapered hollow components, where identifying the location of maximum bending stress is essential for targeted reinforcement.

## Notes

**Competing Interest Statement:** There are no conflicts of interest regarding the publication of this article.

**Funding** This study was supported by Iketani Science and Technology Foundation of Japan.

### Competing Interest Statement

The authors have declared no competing interest.

